# VIVID: a web application for variant interpretation and visualisation in multidimensional analyses

**DOI:** 10.1101/2021.11.16.468904

**Authors:** Swapnil Tichkule, Yoochan Myung, Myo T. Naung, Brendan R. E. Ansell, Andrew J. Guy, Namrata Srivastava, Somya Mehra, Simone M. Caccio, Ivo Mueller, Alyssa E. Barry, Cock van Oosterhout, Bernard Pope, David B. Ascher, Aaron R. Jex

## Abstract

Large-scale comparative genomics- and population genetic studies generate enormous amounts of polymorphism data in the form of DNA variants. Ultimately, the goal of many of these studies is to associate genetic variants to phenotypes or fitness. We introduce VIVID, an interactive, user-friendly web application that integrates a wide range of approaches for encoding genotypic to phenotypic information in any organism or disease, from an individual or population, in three-dimensional (3D) space. It allows mutation mapping and annotation, calculation of interactions and conservation scores, prediction of harmful effects, analysis of diversity and selection, and 3-dimensional (3D) visualisation of genotypic information encoded in Variant Call Format (VCF) on AlphaFold2 protein models. VIVID enables the rapid assessment of genes of interest in the study of adaptive evolution and the genetic load, and it helps prioritising targets for experimental validation. We demonstrate the utility of VIVID by exploring the evolutionary genetics of the parasitic protist *Plasmodium falciparum*, revealing geographic variation in the signature of balancing selection in potential targets of functional antibodies.

## Introduction

The modern explosion of genomics, population genetics and genome-wide association studies (GWAS) is producing enormous amounts of polymorphism data linking nucleotide variation to phenotypic outcomes (Luo, et al. 2011; Uffelmann, et al. 2021) or disease (Duncavage and Tandon 2015; Wu, et al. 2016; Giannopoulou, et al. 2019). Countless informatic and statistical methods have been developed to compile and quantify patterns in genotypic data that can be distilled into biologically meaningful results (Luo, et al. 2011; Uffelmann, et al. 2021). However, visualisation is one of the most powerful methods for pattern recognition because it is closely aligned to our brain’s processing of complex information (Bülthoff and Edelman 1992).

Many tools can display (Variation Viewer; ncbi.nlm.nih.gov/variation/view) and predict the impact of individual single nucleotide polymorphisms (SNPs) on protein sequence and structure for large-scale population comparative studies and GWAS (Glusman, et al. 2017). However, few can display the sum of these SNP locations in protein structures in an automated way. Moreover, current 3D mapping and visualisation tools are either manually-driven (e.g., Chimera; (Pettersen, et al. 2004)) or restricted to specialist applications. cBioPortal (Cerami, et al. 2012), COSMIC-3D (cancer.sanger.ac.uk/cosmic3d/), CRAVAT (Douville, et al. 2013), MoKCa database (Richardson, et al. 2009) and cancer3D (Porta-Pardo, et al. 2015) are limited to cancer-related variants. LS-SNP/PDB (Ryan, et al. 2009), MuPIT (Niknafs, et al. 2013), PopViz (Zhang, et al. 2018) and VarMap (Stephenson, et al. 2019) are limited to humans. SNP2Structure (Wang, et al. 2015) exclusively accesses genetic variation from dbSNP (Sherry, et al. 2001). Finally, to predict the mutational effects of non-cancer and non-human genetic variants on protein structure, users necessarily depend on third-party software and databases. In turn, this requires multiple webservers and input file formats, which is challenging and time-consuming, hampering comparison and accessibility. Furthermore, although existing tools allow annotation of individual SNPs, they do not scale to support the rendering of complex population-level variant data from any given organism or disease.

We present VIVID, a novel interactive and user-friendly platform that automates mapping of genotypic information and population genetic analysis from VCF files in 2D and 3D protein structural space. This platform provides an easy user experience and an integrated analysis environment to yield both individual- and population-level insights, whilst generalising to any organism or disease.

## New Approach

VIVID is a unique ensemble user interface that enables users to explore and interpret the impact of genotypic variation on the phenotypes of secondary and tertiary protein structures. Using data from a standard VCF, VIVID integrates published algorithms, programs, and databases to map, annotate, analyse and visualise the effects of mutations on primary protein sequence, 2D protein residue interactions and a variety of 3D protein-structural renderings at individual and population scales. Fig. 1 provides a schematic workflow for VIVID, which consists of three key components: input, data pre-processing, and visualisation. Overall, VIVID allows the integration of multiple analyses within one interactive visualisation interface, adding new dimensions to variant annotation and functional effect prediction for any organism in any system.

**Figure 1.**
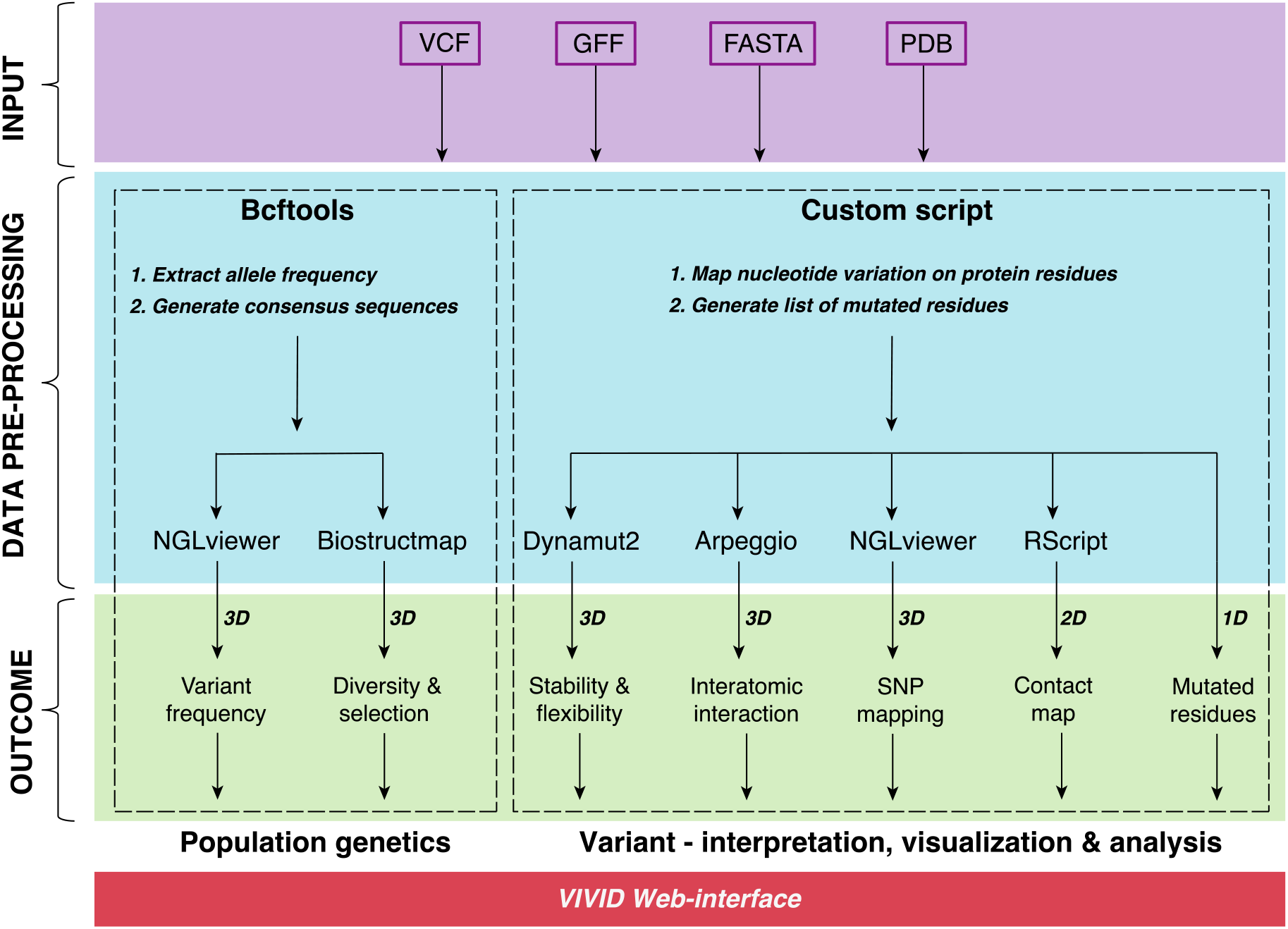
Schematic workflow of VIVID

### Input source

VIVID requires four main inputs in the submission page of the webserver. **1.** The complete nucleotide coding sequence of a gene in FASTA format. **2.** A general feature format (GFF) file to extract genomic coordinates of the queried coding sequence and to map SNPs from **3.** A VCF file on **4.** A protein structure (PDB file), which can either be retrieved on request through VIVID from the AlphaFold2 predicted protein structure database (Jumper, et al. 2021) or provided by users from any source, whether *in silico* predicted (Roy, et al. 2010; Kelley, et al. 2015) or accessed via the RCSB Protein Data Bank API (https://www.rcsb.org)(Berman, et al. 2000). VIVID also allows users to select codon usage preference by selecting appropriate “genetic code” from the drop-down menu (default: standard code).

### Data pre-processing: Mapping mutations in 1D – 3D

SNPs can be identified and explored within the VIVID interface based on their location in 1D linear display, 2D residue contact maps, and 3D protein structural renderings. First, the 1D display is generated by mapping SNPs from the VCF file to the primary protein sequence based on the input GFF file and highlighted in the protein sequence viewer. Next, VIVID uses atomic coordinates from the PDB file to calculate the Euclidian distance between each pair of residues and display each mutated amino acid in the context of long-range residue contacts. Finally, VIVID renders each mutated amino acid in the 3D protein structure using NglViewer (Rose, et al. 2018).

### Data pre-processing: Annotation

Each mutated amino acid is classified as non-synonymous or synonymous using an appropriate genetic code selected by the user. Protein sequences and structural renderings can also link to functional and domain-based annotations using data retrieved from UniProt (The UniProt 2021) and Pfam (Mistry, et al. 2021), if available.

### Data pre-processing: Mutational analyses

VIVID predicts the likely effects of substituted amino acids on protein structure using previously published algorithms, tools and APIs. Dynamut2 (Rodrigues, et al. 2020) is used to predict the effects of missense mutations on protein structure stability and flexibility. Arpeggio (Jubb, et al. 2017) is used to calculate interatomic interactions of mutated residues in 3D space. Amino acid conservation score of mutated residues is computed using the Position-Specific Scoring Matrix (PSSM) of PSI-BLAST (Altschul, et al. 1997).

BCFtools (http://samtools.github.io/bcftools/howtos/install.html) is used to calculate alternative allele frequencies (AF) of each alternate allele from the VCF file. It generates consensus sequences for multiple samples by using a user-supplied coding sequence as a reference, then mapping population genetic observations on protein structures. Finally, a 3D sliding window-based application implemented in BioStructMap (Guy, et al. 2018) is used to map nucleotide diversity (π) and Tajima’s D onto the protein models. This enables users to identify possible links between the effects of natural selection and the structural or biochemical changes in the protein

### Data pre-processing: Visualisation

Nglviewer is used to visualise protein structures in an interactive mode. It also allows the user to map information onto rendered protein structures by selecting options including non-synonymous and synonymous mutations, PSSM conservation scores, nucleotide diversity, Tajima’s D, SNP frequency, etc. In addition, user-selected residues associated with functional and structural domains (retrieved from Uniprot and Pfam databases) in the primary sequence viewer can also be rendered on the protein structure.

## Results

The outcome of VIVID in Fig. 1 are represented in seven panels on the results page of the webserver (Fig. 2).

**Figure 2.**
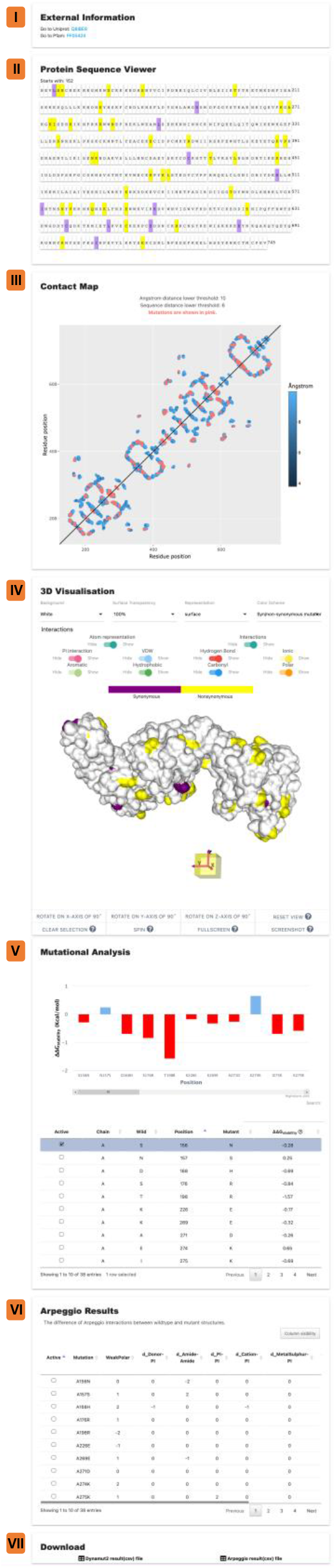
Seven panels showing the VIVID output on the results page. These panels show the results of integrative analyses of SNPs (data derived from the MalariaGen Pf3k version5 dataset) on the EBA175 RII protein.

### External information

The first panel provides an external link for UniProt and Pfam to allow users to render structural and functional domain information (e.g., conserved structural domains, active sites, etc.) on protein sequence and structure to identify mutational hotspots.

### Protein sequence viewer

The second panel displays the translated protein sequence of the residues present in the protein structure where synonymous (purple) and non-synonymous (yellow) mutations are highlighted by default. Users can also click and select protein residues to view them in 3D space in the visualisation panel.

### Contact map

The third panel displays an interactive protein contact map where all pairs of residues mapping within a user-defined Euclidean distance threshold (default: 10 Ångstrom) in 3D-space and found more than six (default) amino acids apart in the primary sequence are displayed. This highlights long-range contacts that are important for stabilising protein structure (Vendruscolo, et al. 1997; Toth-Petroczy, et al. 2016), with pair-wise contacts involving (and potentially disrupted by) mutated residues highlighted in pink.

### 3D visualisation

In the fourth panel, users can visualise the protein structure with different representations and colouring options. Users can map and visualise mutations, annotations and analyses on the 3D model to detect hotspots of genetic variation by selecting various colouring options. Colour coding options include hydrophobicity, non/synonymous mutations, PSSM conservation score, changes of folding free energy (ΔΔ*G*), nucleotide diversity, Tajima’s D, etc.

### Mutational analysis

The fifth panel displays results from mutational analyses performed using Dynamut2 (Rodrigues, et al. 2020). It provides a bar-chart of predicted changes of folding free energy (ΔΔ*G*) of substituted amino acids on protein structure stability and flexibility. The △△*G* values are also reported in tabular format.

### Arpeggio results

The sixth panel reports differences in 20 interatomic interactions between wild-type and mutated residues in tabular format.

### Download

The seventh panel allows users to download Dynamut2 and Arpeggio results.

## Case study

Here, we demonstrate the utility of VIVID and its features by exploring the evolution of vaccine candidates within the parasitic protist *Plasmodium falciparum*, one of the world’s primary causes of malaria (Prugnolle, et al. 2011). Strain-specific immunity hinders vaccine development, and vaccine escape is likely to occur at sites under balancing selection. Drawing on an ensemble of published tools and algorithms, VIVID provides an intuitive and user-friendly interface to analyse functional divergence in more depth. We used SNP data from the published genomes (MalariaGEN Pf3K v5.1) of naturally occurring *P. falciparum* infections from Guinea (n=100) and Thailand (n=148). We filtered these data for SNPs mapping to region II (AA residues: N152–V745) of the Erythrocyte binding antigen protein (EBA175 RII) using the *P. falciparum* 3D7 genome (PlasmoDB release v43). EBA175 mediates binding to human receptor glycophorin A during merozoite invasion of the RBC (Tolia, et al. 2005) (Fig 3A). Region II (RII) of EBA175 is the functional binding domain consisting of 2 cysteine-rich Duffy binding-like domains called F1 and F2 (Sim, et al. 1994). These F1 and F2 domains mediate dimerisation of two EBA-175 proteins through disulphide bond formation (Tolia, et al. 2005), allowing *P. falciparum* merozoites to bind to the erythrocyte surface (Fig 3A) during blood-stage invasion (Tham, et al. 2012). Antibodies that block dimerisation of EBA175 may inhibit glycophorin A binding, which is associated with protection from clinical malaria (Chen, et al. 2013; Irani, et al. 2015) (see residues highlighted in magenta in Fig 3B). Most (> 90 %) of the SNPs mapping to this region in the Guinean (Africa) and Thailand (Asia) populations were non-synonymous (Fig. 3C). Spatially derived Tajima’s D analysis in VIVID identified a large region of the F1 domain (AA residues E226–L294, I312–K324, and W377–I400) of EBA175-RII as being under balancing selection (i.e., positive Tajima’s D values) in both populations (Fig. 3E). Some of the F1 domain mutations, such as E274K, L482V (also under balancing selection), were predicted to stabilise the parasite protein (i.e., positive ΔΔ*G*) (Fig. 3D). Such polymorphisms can be maintained at intermediate frequencies to help the parasite escape host immune responses while maintaining a functional protein. A second region within the F2 domain (AA residues C551–C591 and Y731–F743) was also identified under balancing selection, but this was specific to the Thailand population (circles in Fig. 3E). Some of these residues with the F2 domain are targets of functional antibodies (Ambroggio, et al. 2013; Chen, et al. 2013) (Fig 3B). In contrast, the F1 domain has high nucleotide diversity, less amino acid (low PSSM) conservation and is under balancing selection in both the populations (Fig 3E-G). Therefore, the results suggest geographically variable effectiveness of the EBA175-RII vaccine if the formulation is based on a single reference strain.

**Figure 3.**
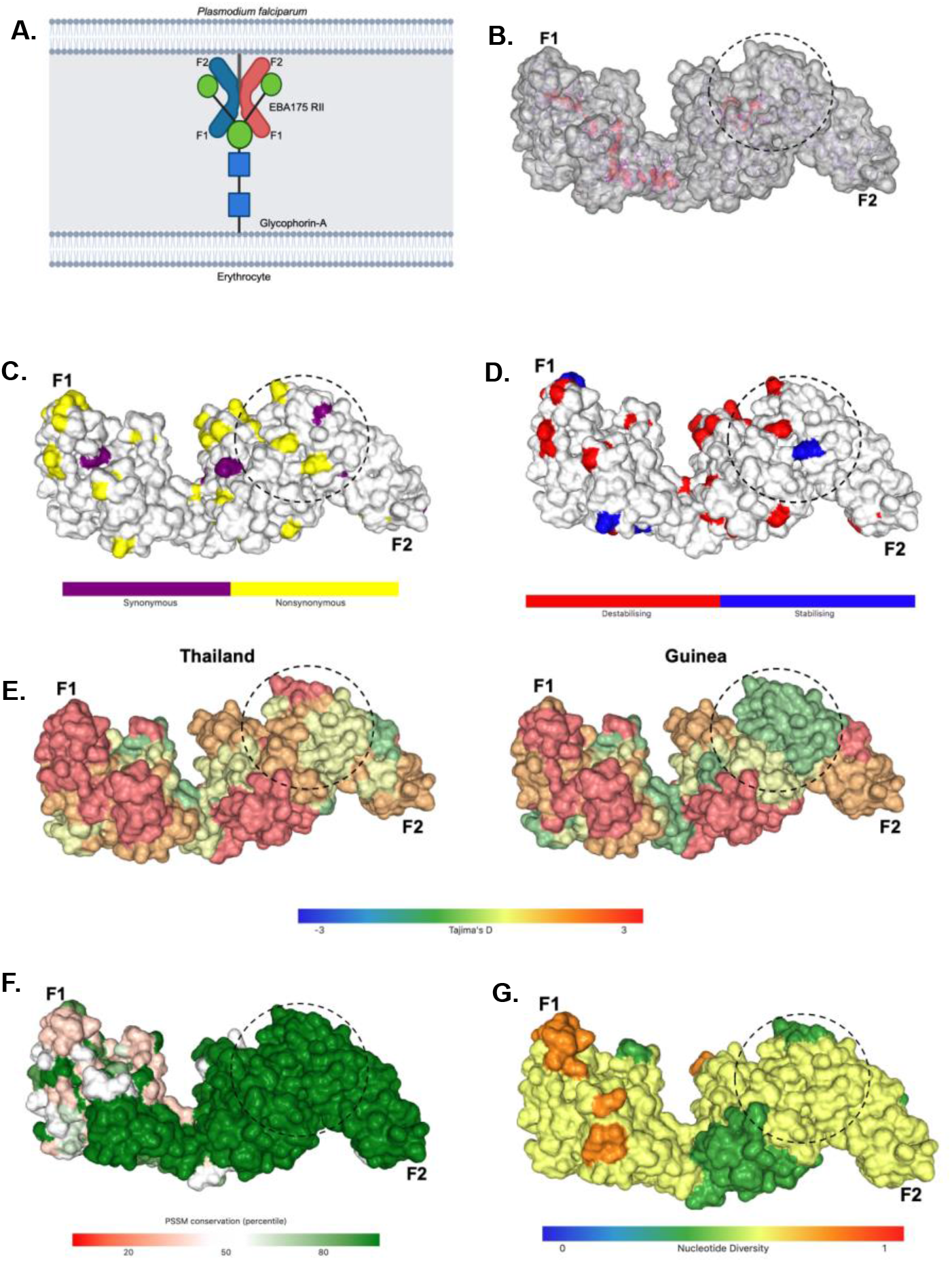
Analyses of EBA175 RII polymorphisms from two different populations - Thailand (Asia) and Guinea (Africa) using VIVID. A part of the F2 domain (circle) illustrates distinct balancing selection between Thailand and Guinea populations. Unless otherwise stated, only the results from the Thailand population are shown. **A**. Schematic diagram of the dimerised EBA175 RII ligand binding to glycophorin A human receptor. **B**. The highlighted residues (magenta) indicated the target of known inhibitory antibodies from previous studies (Ambroggio, et al. 2013; Chen, et al. 2013). **C.** Map of non-synonymous and synonymous mutations on EBA175 RII. **D.** Map of ΔΔG values on EBA175 RII. **E.** Map of spatial Tajima’s D on EBA175 RII. **F**. Map of PSSM scores on EBA175 RII. **G.** Map of nucleotide diversity on EBA175 RII.

## Conclusion

We developed a web-based application, VIVID, to support improved visualisation, analysis and understanding of how mutations impact protein structure and function in any organism, using a selection of standardised input format files. This method allows users to examine mutations from individuals or populations in multiple dimensions for solved or *in silico* predicted protein structures, which can either be imported directly from RCSB or AlphaFold2, or uploaded by the user. VIVID can assess SNPs for their impact on protein structure, inferred function, and surface biochemistry (e.g., hydrophobicity) using individual-based data. For population studies, in addition to these features, VIVID can use the SNP data to colour any protein model based on localised differences in population genetic metrics. This enables users to visualise protein evolution in 3D and identify genomic regions under selection. The architecture of this browser-based tool is designed to benefit the broader scientific community, provide an easy user experience, and offer an integrated analysis environment. Our web portal integrates features from multiple programs, algorithms and databases, providing users with a single platform to explore and interpret variants in multiple dimensions. In addition, the user-friendly and interactive panels allow downloading multiple result outputs that can be used downstream in other programs. In this paper, we have demonstrated the utility of VIVID by using a case study of the evolution of vaccine candidates against malaria. This will be a valuable resource to the research community working in structural and functional proteomics and genomics and will have numerous valuable applications.

## Future developments

We are in the process of expanding the usability of this application by integrating additional features from the available wealth of bioinformatics resources. These include widely used variant repositories for model-organisms, annotation databases in addition to UniProt, and additional protein functional properties to map on 3D protein structures. Moreover, we plan to integrate other developed algorithms and tools from our Biosig server (http://biosig.unimelb.edu.au/biosig). Furthermore, we are in the process of expanding VIVID features for population and evolutionary genomics analyses and 3D visualisation. Finally, we will make available a standalone package for free download. In addition to the current analyses, this will allow users with sufficient computational resources to scale VIVID to perform proteome-wide structural studies.

## Code and data availability

The server is publicly available at http://biosig.unimelb.edu.au/vivid. The SNP dataset used in the case study was obtained from (MalariaGEN Pf3K (version 5.1), https://www.malariagen.net/data/pf3k-5)

## Funding source

This work was supported by the Australian National Health and Medical Research Council (APP1126395 and GNT1174405), Victorian State Government Operational Infrastructure Support and Australian Government National Health and Medical Research Council Independent Research Institute Infrastructure Support Scheme to A. R. J. and D.B.A. This work was supported by a Victorian Health and Medical Research Fellowship from the Victorian State Government to B.P. This work was supported by an NHRMC Principal Research Fellowship (GNT1155075) to I.M. A.G was supported by a Vice-Chancellor’s Postdoctoral Research Fellowship from RMIT University. S.T. was supported by a Walter and Eliza Hall International PhD Scholarship. S.T., Y.M. and M.T.N were further supported by a Melbourne Research Scholarship.

## Acknowledgments

S.T. acknowledges the Australian Society for Parasitology (ASP) for a student conference travel grant and a JD Smyth Postgraduate Travel Award for a Network Researcher Exchange, Training and Travel Award. In addition, S.T. acknowledges the IT team at the Walter and Eliza Hall Research Institute, Australia.

## Competing interests

The authors declare that there are no conflicts of interest.

## Notes

https://www.malariagen.net/data/pf3k-5

### Competing Interest Statement

The authors have declared no competing interest.

